# Rap1, AF6 and Pak4 cooperate with Par3 to promote *ZA* assembly and remodeling in the fly photoreceptor

**DOI:** 10.1101/155945

**Authors:** Rhian F. Walther, Mubarik Burki, Noelia Pinal, Clare Rogerson, Franck Pichaud

## Abstract

The epithelial *Zonula adherens (ZA)* is a main adhesion compartment that enables organogenesis by allowing epithelial cells to assemble into sheets. How *ZA* assembly is regulated during epithelial cell morphogenesis is not fully understood. We show that during *ZA* morphogenesis, the function of the small GTPase Rap1 and the F-actin binding protein AF6/Cno are both linked to that of the P21-activated kinase Pak4/Mbt. We find that Rap1 and Mbt regulate each other’s localization at the *ZA* and cooperate in promoting ECadherin stabilization. During this process Cno regulates the recruitment of Baz at the *ZA,* a process that is also regulated by Arm phosphorylation by Mbt. Altogether, we propose that Rap1, Cno and Mbt regulate *ZA* morphogenesis by coordinating ECadherin stabilization and Baz recruitment and retention. In addition, our work uncovers a new link between two main oncogenes, Rap1 and Pak4/Mbt, in a model developing epithelial cell.

## Introduction

The epithelial *ZA* consists of a lateral circumferential belt of *Adherens Junction* (*AJ*) material that allows for epithelial cells to assemble into sheets. Loss of epithelial adhesion is a hallmark of cancer and there is therefore a strong interest in better understanding how *ZA* morphogenesis, remodeling and maintenance are regulated. The adhesion molecule ECadherin/Shotgun (ECad) and its effector βcatenin/Armadillo (Arm) are main *AJ* components of the *ZA* in animal epithelial cells. Work in *Drosophila* and vertebrate cells points to multiple pathways that regulate *AJ* material morphogenesis during epithelial morphogenesis, including membrane delivery, endocytosis, local accumulation and stabilization at the plasma membrane (Bryant and Stow, 2004; Tepass, 2012). However, we still lack an integrated view of how these pathways and the corresponding molecular players come together to regulate *ZA* morphogenesis and remodeling during development.

The fly retina has long been used as a model system to study the genetic and molecular basis of *ZA* morphogenesis and remodeling during organogenesis. During pupal development, photoreceptors build a new *ZA* to accommodate the nascent apical light-gathering structure, called the rhabdomere (Ready, 2002). During this process the Par complex, which consists of Cdc42-Par6-aPKC and Par3/Bazooka (Baz), regulates the specification of the photoreceptor cortex and plasma membrane into a sub-apical domain (stalk membrane) and *ZA* (Hong et al., 2003; Nam and Choi, 2003; Walther et al., 2016; Walther and Pichaud, 2010). During *ZA* morphogenesis, phosphorylation of Baz by aPKC at the conserved S980 leads to the apical exclusion of P-S980-Baz, a step of molecular sorting that also depends on Crumbs (Crb) capturing Par6-aPKC and Stardust (Sdt) at the stalk membrane (Krahn et al., 2010; Morais-de-Sa et al., 2010; Walther and Pichaud, 2010). Confined to the apical-lateral border of the cell, P-S980-Baz is thought to promote *ZA* assembly by binding to Arm and Echinoid (Ed), two main *AJ* components (Wei et al., 2005).

Next to Baz, the Cdc42 effector type-2 p21-activated kinase Mushroom bodies tiny (Mbt/Pak4) has also been shown to regulate *ZA* morphogenesis in several epithelial cell types by promoting Baz retention at the *ZA* and regulating the accumulation of the ECad-Arm complex via phosphorylating βCat/Arm (Jin et al., 2015; Law and Sargent, 2014; Menzel et al., 2008 Schneeberger, 2003 #1892; Walther et al., 2016). In vertebrate cells, Pak4 has also been linked to the Par complex via Par6b phosphorylation, indicating a potential cross talk between Par6b-aPKCγ/ζ and Pak4, downstream of Cdc42 (Jin et al., 2015; Wallace et al., 2010). While in fly photoreceptors both Mbt and Baz are main regulators of *AJ* accumulation at the developing *ZA,* they seem to operate as part of parallel convergent pathways. This is demonstrated by the fact that while Arm accumulation is reduced in *mbt* null mutant photoreceptors, *AJ* material is no longer present at the plasma membrane in cells mutant for both *mbt* and *baz* (Walther et al., 2016). How exactly Baz contributes to regulating *AJ* material accumulation at the *ZA* is not fully understood. Similarly, phosphorylation of βCat/Arm by Pak4/Mbt (Law and Sargent, 2014; Menzel et al., 2008) cannot fully account for *mbt* function during *ZA* morphogenesis, as re-introducing a phosphomimetic form of Arm does not rescue the loss of *mbt* function (Walther et al., 2016). Therefore other factors must regulate *AJ* morphogenesis during *ZA* maturation, either in concert with Mbt and Baz, or as part of the Mbt and Baz pathways.

An interesting candidate in contributing to *AJ* accumulation at the epithelial *ZA* is *Rap1*. This member of the Ras subfamily of small GTPases has been shown to localize at the *AJ* in various fly epithelia, and to be an essential *AJ* regulator (Boettner et al., 2003; Boettner and Van Aelst, 2007; Choi et al., 2013; Knox and Brown, 2002; O’Keefe et al., 2009; Spahn et al., 2012; Wang et al., 2013). In the early embryo, Rap1 and its effector F-actin binding protein AF6/Cno (Boettner et al., 2003; Mandai et al., 2013; Sawyer et al., 2009), regulate the apical localization of both Baz and Arm, with Baz reciprocally influencing Cno localization. Later in embryonic development, Baz is required at the *ZA* to capture preassembled *AJ* material, thus promoting *ZA* morphogenesis (McGill et al., 2009). In addition, work in human MCF7 cells has shown a role for Rap1 during *AJ* maturation via promoting ECad recruitment at the sites of cell-cell contact, a function that has been shown to be mediated, at least in part, by Cdc42 (Hogan et al., 2004). Here we sought to examine the relationship between Rap1, its GEF Dizzy/PDZ-GEF and the protein network that drives epithelial apical cortex and plasma membrane specification using the pupal photoreceptor as a model system.

## Results

### Dizzy and Rap1 regulate pupal photoreceptor *ZA* morphogenesis

In the fly retina, Rap1 has been previously shown to regulate *AJ* remodeling between newly specified photoreceptors, and between retinal accessory cells (cone and pigment cells) (O’Keefe et al., 2009). To test whether *dizzy* and *Rap1* are required during *ZA* morphogenesis in the pupal photoreceptor, we made use of the *rap1-Rap1::GFP* and *dizzy-Dizzy::GFP* transgenes, which allow for expression of these proteins under their endogenous promoter. We found that Rap1::GFP accumulates predominantly at the developing pupal photoreceptor *ZA* (Figure 1A), and can also be detected at lower levels at the apical photoreceptor membrane, which includes the stalk membrane. Dizzy::GFP (Figure 1B’) shows a similar pattern, with perhaps less accumulation at the apical membrane when compared to Rap1::GFP. Accumulation at the *ZA* and low levels of expression at the apical membrane is also observed for Arm (Walther and Pichaud, 2010); and (Figure 1A”, 1B). Therefore in the developing pupal photoreceptor, the expression pattern of Dizzy, Rap1 and Arm are very similar.

**Figure 1:**
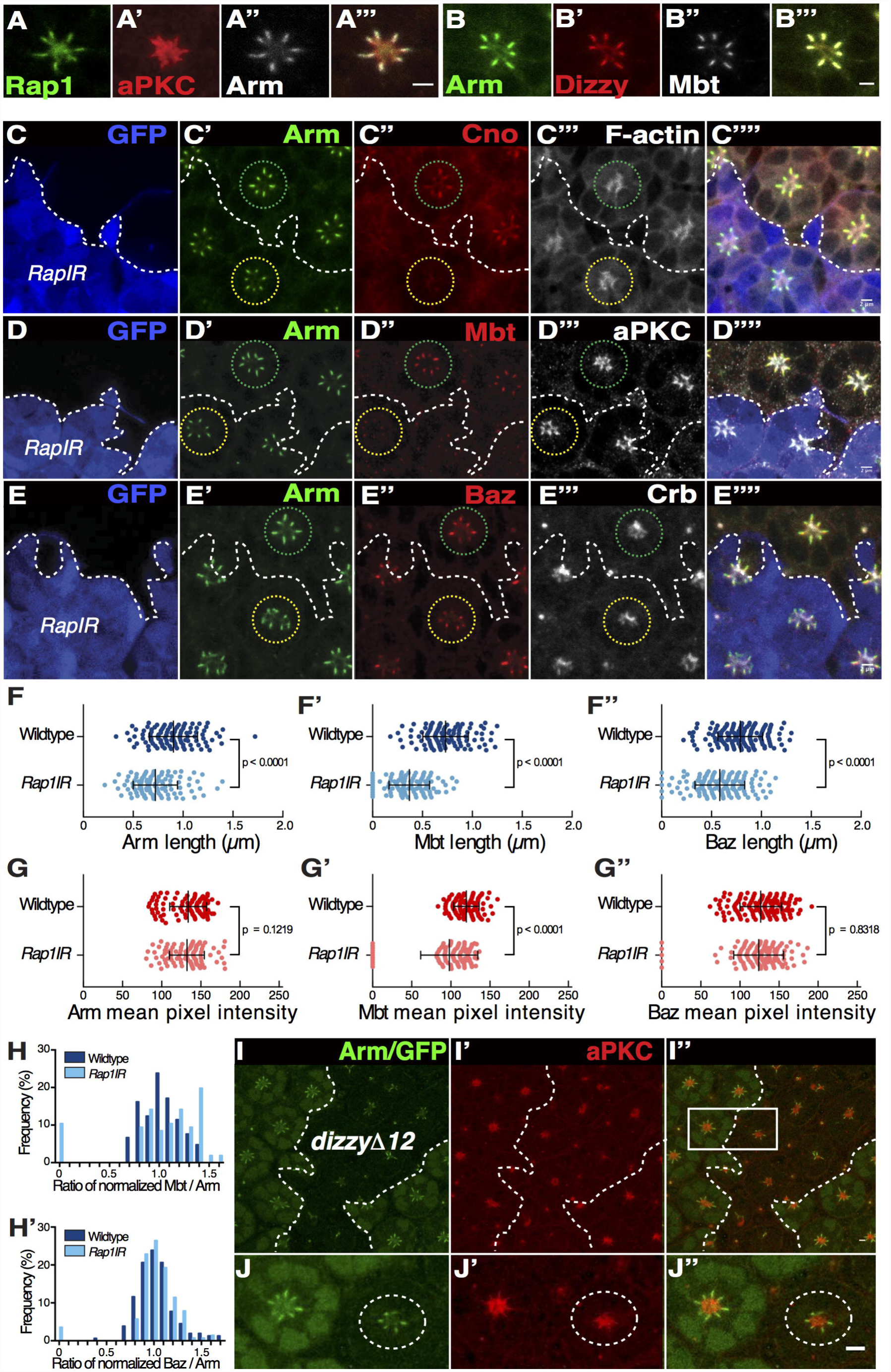
Dizzy and Rap1 regulate the accumulation of *AJ* material during *ZA* morphogenesis. (A-A”’) Rap1::GFP (A), aPKC (A’) and Arm (A”). Scale bar 2μm. (B-B”’) Arm (B), Dizzy::GFP (B’) and Mbt (B”); scale bar = 1.5μm. (C-E) *Rap1IR* cells positively labeled by nuclear GFP (blue) and stained for Arm (C’, D’, E’), Cno (C”), Mbt (D’’), Baz (E”) and F-actin (C”’), aPKC (D”’) and Crb (E”’). Scale bar = 2μm. (F-F”) Quantification of Arm (F), Mbt (F’) and Baz (F”) domain length at the *ZA*. (G-G”) Quantification of Arm (G), Mbt (G’) and Baz (G”) domain mean pixel intensity at the *ZA*. (H) Ratio of normalized Mbt and Arm mean pixel intensity. (H’) Ratio of normalized Baz and Arm mean pixel intensity. (I-I”) *dizzydΔ^12^* mutant clone labeled by the lack of nuclear GFP staining (I), Arm (I and J), aPKC (I’ and J’). A dashed line highlights the contour of the *dizzydΔ^12^* mutant clone. Scale bar = 20μm. A square box highlights two contiguous ommatidia: one wild type and one mutant. (J-J”) close up of the two ommatidia highlighted in (I”). A dashed line circles the *dizzydΔ^12^* mutant ommatidium. Scale bar = 2μm.

*Rap1* is required to preserve the integrity of the retina (Supplementary Figure 1). Generating mutant clones using the strong allele *Rap1^CD3^*, or expressing high levels of a previously validated *Rap1IR* construct (O’Keefe, 2009 #1401), leads to severe defects in recruiting the full complement of retinal accessory cells including the cone cells (Supplementary Figure 1A). Missing cone and pigment cells lead to retinal cell delamination, with many photoreceptors found below the floor of the retina (Supplementary Figure 1B-D), preventing us from assessing polarity and *ZA* morphogenesis. In order to bypass this strong phenotype we limited the expression of *Rap1IR* to the pupal phase. Decreasing the expression of *Rap1* at pupal stages did not affect photoreceptor apical-basal polarity in the majority of ommatidia examined (Figure 1C-E). However, we could measure shorter *ZA* (as measured along the apical-basal axis of the cell), and thus an overall decrease in the quantity of Arm at the *ZA* of the *Rap1* deficient photoreceptors (Figure 1C’, D’, E’ and 1F). A survey of *ZA*-associated proteins revealed that in addition to Arm, the length of the Mbt (Figure 1D” and 1F’) and Baz (Figure 1E” and 1F”) domains were also significantly reduced in *Rap1IR* photoreceptors. Furthermore, Cno could no longer be detected at the *ZA* (Figure 1C”) and Mbt levels were significantly decreased when compared to wild type (Figure 1D” and 1G’). In some cases, *ZA* domains were present that did not contain Mbt, resulting in a significant alteration of the ratio between Mbt and Arm. In wild type, *ZA* levels of Mbt/Arm are correlated and follow a normal distribution. In *Rap1IR ZA* this correlation was disrupted, with significantly more junctions presenting either high or low Mbt/Arm ratios that fall outside of a normal Gaussian distribution (Figure 1H). In these shortened *ZA*, levels of Arm and Baz were comparable to wild type (Figure 1G, 1G”) and the ratios of Baz/Arm in *Rap1IR* cells remained similar to wild type (Figure 1H’). Apical levels of F-actin (Figure 1C”’), aPKC (Figure 1D”’), and Crb (Figure 1E”’), were not affected in *Rap1IR* photoreceptors when compared to wild type. These data indicate that Rap1 is required for the accumulation of *AJ* material at the developing *ZA*. This function appears most critical when considering the *ZA* levels of Cno and Mbt,

Next, to examine the function of *dizzy* during *ZA* morphogenesis, we made use of the strong *dizzy^Δ12^* allele. We found that reducing *dizzy* expression leads to a phenotype similar to that seen in *Rap1IR* photoreceptors, including a shortening of the *ZA* along the apical-basal axis (Figure 1I-J). Consistent with Dizzy acting as a Rap1-GEF in photoreceptors, removing a copy of the *dizzy* locus enhances the mild rough-eye phenotype obtained when reducing the expression of *Rap1* using RNAi (Supplementary Figure 2A-D).

### *Rap1* promotes *AJ* stabilization during *ZA* remodeling

We have previously shown that in pupal photoreceptors, loss of *mbt* function leads to an increase in the mobile fraction of ECad at the *ZA* when compared to wild type over 250 secondes (Walther et al., 2016). Our analysis of *Rap1IR* indicates that Mbt accumulation is reduced in the corresponding *ZA* (Figure 1D” and 1G’), which should therefore be accompanied by an increased in ECad mobility. To assess whether this is the case, we made use of FRAP and compared the recovery after photo-bleaching of a *ubi-ECad::GFP* transgene in wild type and *Rap1IR* photoreceptors. In wild type cells, over approximately 250 sec, we estimated that 25% of ECad::GFP is mobile, which is consistent with previous estimations from our lab (Walther et al., 2016) (not shown).

However, while ECad::GFP shows a stronger recovery over this relatively short time scale in *Rap1IR* when compared to wild type, the GFP signal failed to plateau (not shown), preventing us from extrapolating the mobile fraction. We therefore performed FRAP over a longer time scale (1000 sec). Over this long time scale, we found approximately 35% of ECad::GFP is mobile in wild type *ZA*, while ~70% is mobile in *Rap1IR* photoreceptors (Figure 2A-C). These data indicate that Rap1 promotes ECad stabilization at the *ZA,* and are compatible with Mbt mediating part of Rap1 function during this process.

**Figure 2:**
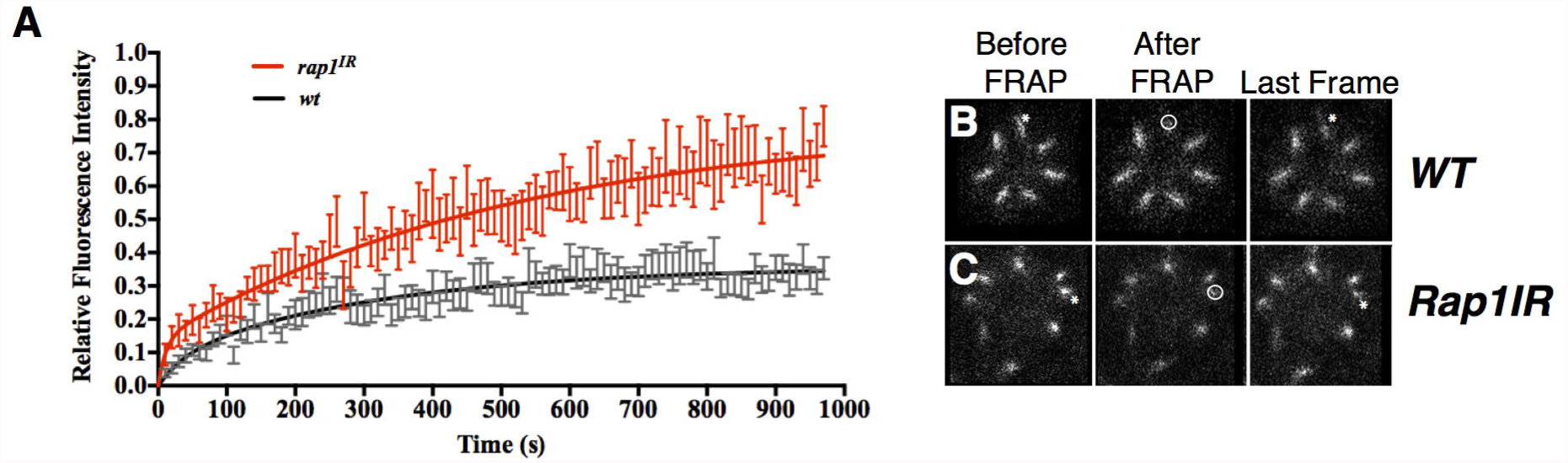
Rap1 regulates the stability of ECad at the developing *ZA*. (A) FRAP fit for ECad::GFP in wild type (black) and *Rap1IR* (red) photoreceptors. For both genotypes, the basal end of the developing *ZA* (dashed circle) was photo-bleached (B-C). For wild type *ZA* FRAP, n = 14 and for *RapIR,* n = 12.

### Cno couples Arm and Baz at the *ZA* and is required for the apical accumulation of aPKC and Crb

Next to regulating Mbt accumulation at the *ZA*, one likely mechanism whereby Rap1 might promote ECad stabilization is through the F-actin linker Cno (Kooistra et al., 2007). In the pupal photoreceptor, Cno localizes at the *ZA* and this localization is also strongly decreased in *Rap1IR* photoreceptors (Figure 1C”). Decreasing *cno* expression using the strong *cno^R2^* allele leads to delamination of the mutant photoreceptors through the floor of the retina (Figure 3A-B), a phenotype resembling that obtained when strongly reducing *Rap1* expression. As for *Rap1* loss of function, delamination of the *cno^R2^* mutant retinal cells is likely due to strong defects in assembling the full complement of interommatidial accessory cells, and polarity of the delaminated photoreceptors is strongly compromised (Figure 3B). The delamination phenotype complicates the analysis of *cno* function. In order to circumvent this issue we made use of *cno* RNAi (*cnoIR*). Examining retinas mosaic for *cnoIR* revealed that this factor is required for the accumulation of Arm (Figure 3C’, 3E’), Baz (Figure 3C”) and Mbt (Figure 3E”) at the developing photoreceptor *ZA*. Examining *cnoIR* mutant *ZA*, we noted instances where Arm was present at the *ZA* but Baz or Mbt were absent (Figure 3D, 3F). The change in the relative accumulation of Arm and Mbt measured in *cnoIR* resembles that quantified in *Rap1IR* (Figure 1H), suggesting that Rap1 and Cno function during *ZA* morphogenesis are linked. However, in the case of *cnoIR*, we also detect uncoupling between Arm and Baz, a phenotype not detected in *Rap1IR*, indicating that part of Cno function is independent of Rap1. In addition, levels of Crb and aPKC were decreased in *cnoIR* mutant cells (Figure 3C”’ and 3E”’), indicating a Cno regulates the accumulation of these factors during apical membrane morphogenesis.

**Figure 3:**
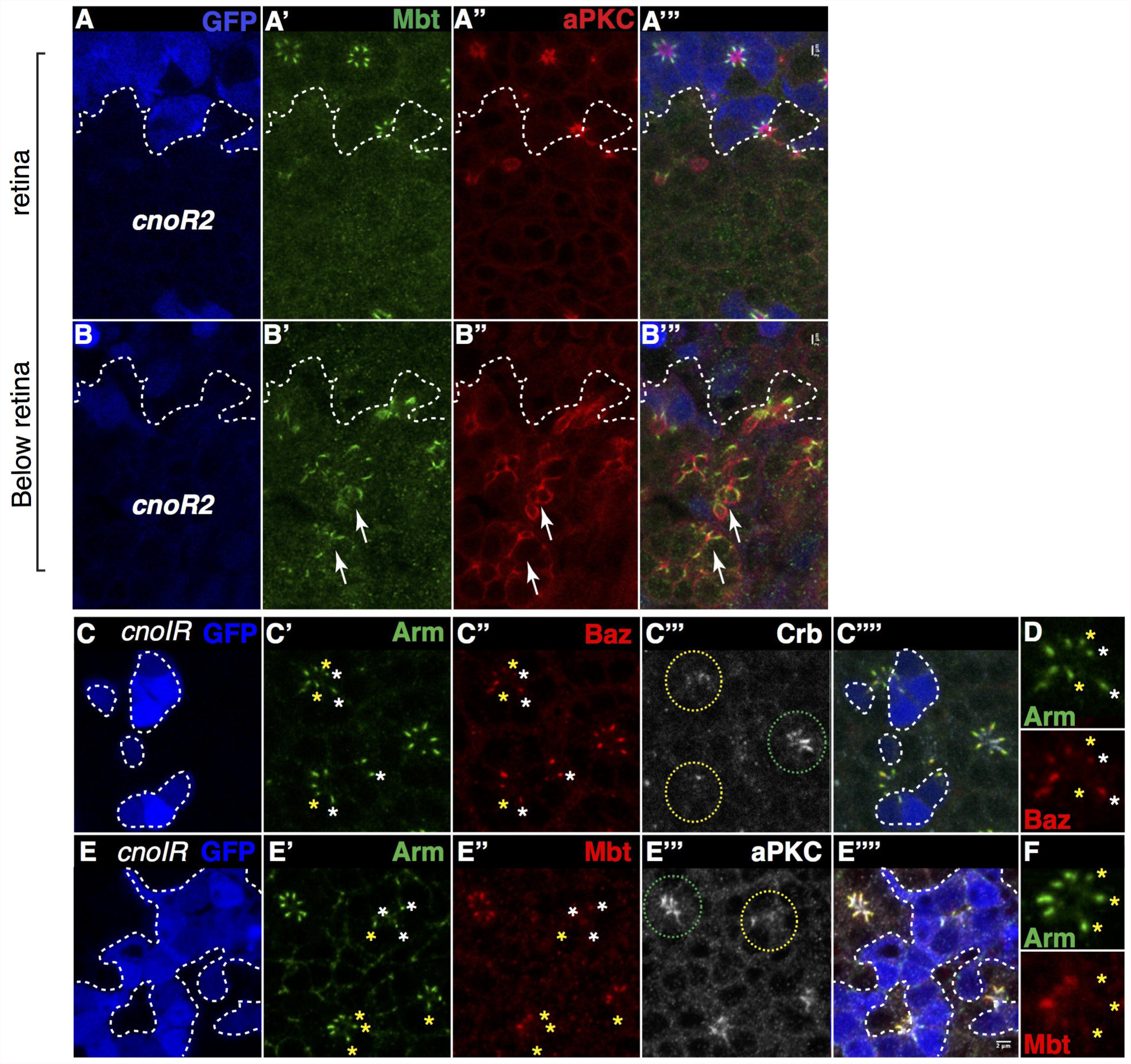
Cno regulates the coupling of Arm, Baz and Mbt at the developing *ZA*. (A-B) *cno^R2^* mutant cells labeled by the lack of nuclear GFP (blue) stained for Mbt (A’ and B’) and aPKC (A” and B”). White arrows indicate *cno^R2^* mutant photoreceptors that have delaminated from the retinal neuroepithelium. (C-F) *cnoIR* clones labeled by a nuclear GFP (C and E) and stained for Arm (C’ and E’), Baz (C”), Crb (C”’), Mbt (E”) and aPKC (E”’). (D and F) show a magnification of one mosaic ommatidium to highlight the absence of Baz (D) or Mbt (F) in some of the Arm domains. White stars label *ZA* containing both Arm and Baz, while yellow stars indicate *ZA* containing Arm but depleted for Baz (D) or Arm but depleted for Mbt (F). Scale bars = 2μm.

### Linking *Rap1* function to that of *baz* and *mbt*

Our results suggest that during *ZA* morphogenesis, Rap1 could function through Cno, and Mbt. In addition, Cno accumulation at the *ZA* depends on *mbt* (Figure 4A”), indicating multiple cross talks exist between Rap1, Cno and Mbt. To examine the relationship between Rap1, Cno and Mbt, we asked whether expressing Mbt or Cno could ameliorate the *Rap1IR ZA* phenotype. Expressing *mbt* in *Rap1IR* cells did not ameliorate the length of the *ZA* and did not restore levels of Cno (Figure 4B”, 4D). Expressing *cno* in *Rap1IR* cells did not ameliorate the length of the *ZA* (Figure 4C-D). These results indicate that Rap1 function on *ZA* morphogenesis is pleiotropic.

**Figure 4:**
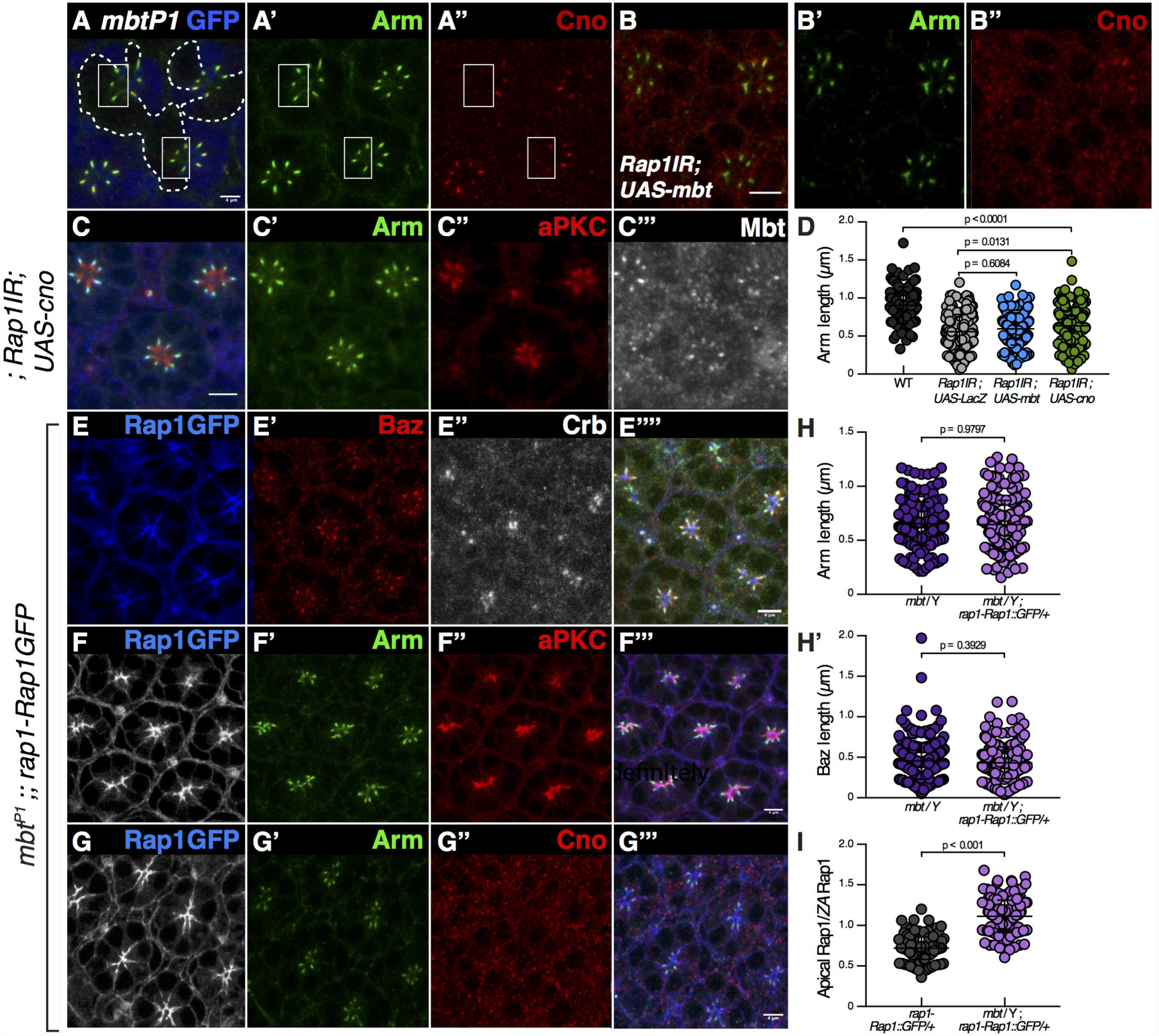
Rap1 and Mbt synergize during photoreceptor *ZA* morphogenesis. (A) *mbt^P1^* mutant photoreceptors labeled by the lack of the nuclear GFP (blue) and stained for Arm (A’) and Cno (A’’). (B) *Rap1IR* photoreceptors expressing *mbt* and labeled for Arm (B’) and Cno (B’’). (C) *Rap1IR* photoreceptors expressing *cno* and labeled for Arm (C’), aPKC (C’’) and Mbt (C’’’). (D) Quantification of *ZA* length in wild type (black), *Rap1IR; UAS-LacZ* (grey) *Rap1IR*; *UAS-mbt*, (blue) *Rap1IR*; *UAS-cno* (green). (E-G) *mbt^P1^* mutant photoreceptors expressing *rap1-Rap1::GFP* (E, F, G) stained for Baz (E’), Arm (F’, G’), Crb (E’’), aPKC (F’’) and Cno (G’’). (H) Quantification of the length of the Arm and (H’) Baz domains at the *ZA* in *mbt^P1^* mutant and *mbt^P1^* mutant expressing *rap1-Rap1::GFP.* (I) Ratios between apical Rap1::GFP signal and Rap1::GFP signal associated to the *ZA.* Scale bars = 2μm.

Next, to examine the relationship between *Rap1* and *mbt* in more detail, we asked whether expressing the *rap1-Rap1::GFP* transgene could ameliorate the decrease in *AJ* material accumulation measured in *mbt^P1^* null mutant photoreceptors. *mbt^P1^* mutant cells are characterized by a decreased accumulation of Arm, Baz (Walther et al., 2016) (Supplementary Figure 3A-B), and Cno (Figure 4A”) at their *ZA*. When expressing *rap1-Rap1::GFP* in *mbt^P1^* mutant cells (Figure 4E-G), we did not measure any significant recovery in the length of the Arm (Figure 4F’, G’ and 4H) or Baz domains (Figure 4E’, 4H’) when compared to *mbt^P1^* mutant cells, and Cno levels were not restored (Figure 4G”). However, we noted that Rap1::GFP expression was more widespread than in wild type photoreceptors, as the preferential *ZA* accumulation (Figure 1A) was no longer readily detected. Instead Rap1::GFP was localized all over the apical membrane (Figure 4E, 4F, 4G and 4I). These results indicate that Mbt regulates the distribution of Rap1 between the apical membrane and the *ZA*.

Finally, we made use of genetics to probe the relationship between *Rap1* and *baz*. Firstly, we found that *Rap1* and *baz* genetically interact during eye development, as decreasing the expression of *baz* using RNAi (*bazIR*), enhances the *Rap1IR* rough eye phenotype (Supplementary Figure 2A-B, 2E-F). Secondly, to assay whether *Rap1* function during *ZA* morphogenesis relates to that of Baz we generated photoreceptors deficient for both *baz* (using the *baz^xi106^* allele) and *Rap1* (using the *NP-Gal4^2631^*-*Rap1IR* strain) (O’Keefe et al., 2009). As we have show before (Walther et al., 2016), *AJ* material such as Arm is detected at the plasma membrane in *baz^xi106^* and *mbt^P1^* single mutant cells (Figure 5A and Supplementary Figure 3). However, no *AJ* material is detected in *baz^xi106^*, *mbt^P1^* double mutant cells (Figure 5B) indicating that *baz* and *mbt* converge in promoting *AJ* material accumulation at the plasma membrane. We found that expressing *Rap1IR* in *baz^xi106^* photoreceptors led to fewer cortical domains positive for Arm shared by flanking photoreceptors when compared to *baz^xi106^* and *Rap1IR* single mutant cells (Figure 5C, 5E). This was accompanied by a loss of Mbt accumulation (Figure 5C”’), which is consistent with our observation that *Rap1* is required for the accumulation of Mbt at the *ZA* (Figure 1D”; quantified in 1F’ and 1G’). These data further indicate that during *ZA* morphogenesis, the function of *Rap1* and *mbt* are interlinked.

**Figure 5:**
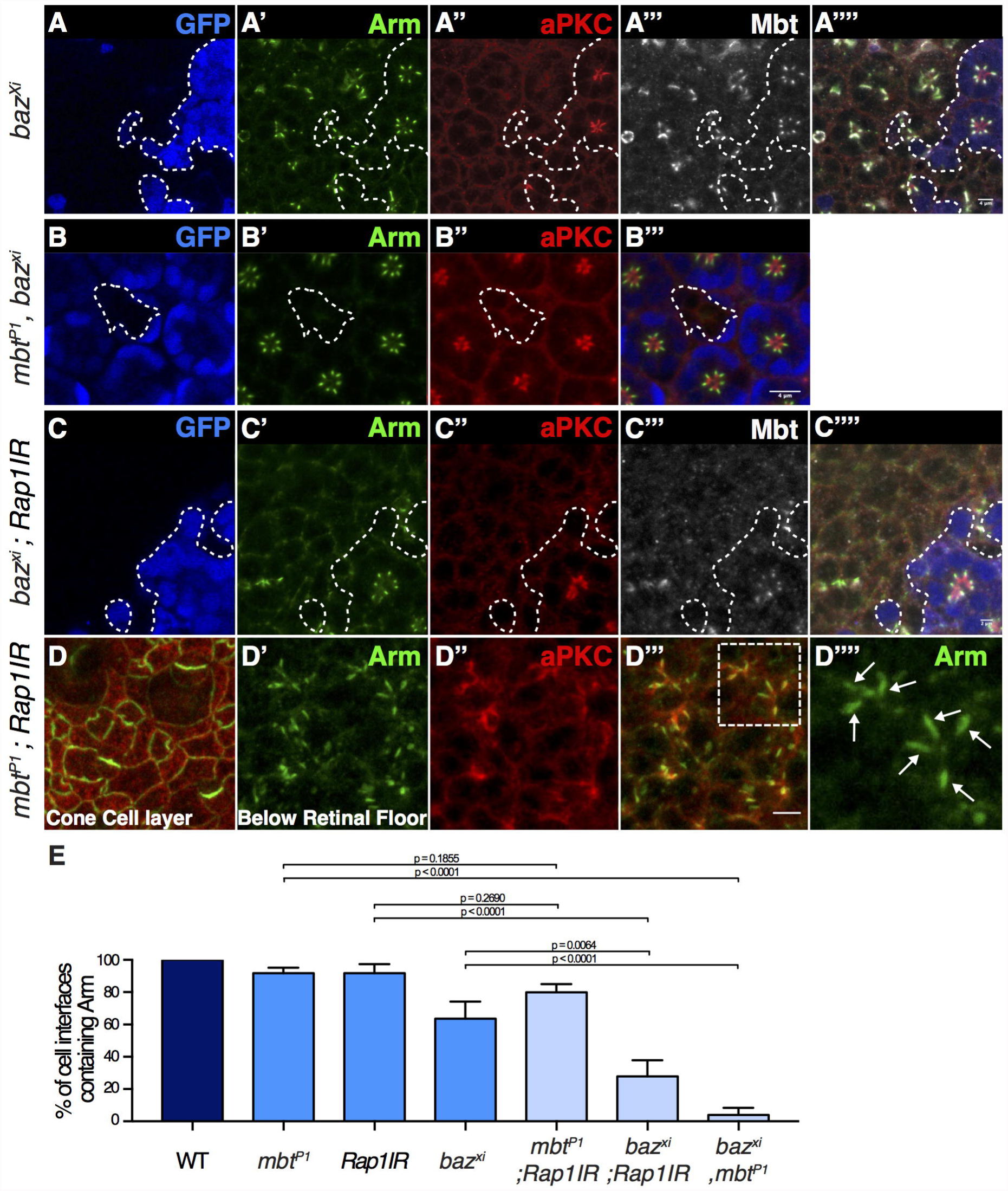
Rap1 and Mbt function together during *ZA* morphogenesis. (AA’’’’) *baz^xi106^* mutant cells labeled by the lack of nuclear GFP (blue) and stained for Arm (A’), aPKC (A’’) and Mbt (A’’’). (B-B’’’) *mbt^P1^*, *baz^xi106^* double mutant cells labeled by the lack of nuclear GFP (blue) and stained for Arm (B’) and aPKC (B’’). (C-C’’’’) *baz^xi106^, Rap1IR* double mutant cells are labeled by the lack of nuclear GFP (blue) and stained for Arm (C’), aPKC (C’’) and Mbt (C’’’). (D) Confocal section of the cone and pigment cells in an *mbt^P1^; Rap1IR* retina stained for Arm (green) and aPKC (red). (D’-D’’’’) View of the delaminated photoreceptor proximal to (D). (D’) Arm, (D’’) aPKC, (D’’’) Merge (D’’-D’’’); a white-dashed rectangle highlights 2 ommatidia that are magnified in (D””). White arrows point to *ZA* domains between flanking photoreceptors. (E) Quantification of the % of pairs of photoreceptors sharing a lateral Arm domain in wild type, *mbt^P1^, Rap1IR, baz^xi106^,* double *mbt^P1^; Rap1IR,* double *baz^xi106^; Rap1IR* and double *baz^xi106^, mbt^P1^*. Scale bars = 4μm.

To complement these experiments, we next asked whether decreasing *Rap1* expression could modify the *mbt* phenotype. Combining the null allele *mbt^P1^* to *Rap1IR* led to a very strong additive effect as nearly all photoreceptors delaminated from the retina, a phenotype due to strong defects in recruiting cone and pigment cells around the photoreceptor clusters (Figure 5D). Nevertheless, a majority of the delaminated, photoreceptors still presented Arm domain linking flanking photoreceptors (Figure 5D”” and 5E). Altogether, these genetic experiments argue in favor of *Rap1* functioning together with *mbt* during Baz-dependent *ZA* morphogenesis.

## Discussion

In the pupal photoreceptor, *ZA* morphogenesis is orchestrated by a conserved protein network that includes the Par complex, Crb and its binding partners Sdt and PATJ, together with the lateral kinase Par1 (Berger et al., 2007; Hong et al., 2003; Izaddoost et al., 2002; Nam and Choi, 2003; Pellikka et al., 2002; Richard et al., 2006; Walther et al., 2016; Walther and Pichaud, 2010). While loss of *ZA,* for example in *arm^3^* mutant photoreceptors, leads to complete failure in pupal photoreceptor apical-basal polarization (Walther et al., 2016), the connection between the *ZA* and the protein network that drives its morphogenesis is not fully understood. We and others have previously shown that Mbt regulates pupal photoreceptor development by promoting *ZA* morphogenesis (Menzel et al., 2007; Walther et al., 2016). During this process Mbt contributes in preventing Baz from spreading to the lateral membrane, a regulation we have found depends on the phosphorylation of Arm by Mbt at S561 and S688 (Walther et al., 2016).

Our results show that Mbt function is linked to that of Rap1 and Cno. We find that Cno couples Arm and Baz accumulation at the *ZA*, as we detect *ZA* domains that do not contain Baz in *cnoIR* photoreceptors. This phenotype resembles that seen when substituting Arm by an Mbt-phospho-dead version of Arm (Walther et al., 2016). Such uncoupling between Arm and Baz is not detected in *Rap1IR* photoreceptors. While this indicates that part of *cno* function is independent of *Rap1*, it suggests that Mbt and Cno function during photoreceptor morphogenesis are linked. Our work shows that Cno is nearly absent at the *ZA* of *mbt* mutant photoreceptors. We therefore propose that part of Mbt’s function in promoting *ZA* accumulation of Baz is mediated by Cno. Whether Cno accumulation at the photoreceptor *ZA* is linked to Arm phosphorylation at S561 and S688 is an interesting possibility that remains to be investigated. *mbt* mutant photoreceptors are frequently found below the floor of the retina, and unlike for *cno* and *Rap1*, this phenotype is not due to defects in retinal accessory cells, but instead is autonomous to the photoreceptor (Walther et al., 2016). These *mbt* photoreceptors present strong defects in apical-basal polarity when considering aPKC and Crb for example (Walther et al., 2016). We show here that *cnoIR* photoreceptors can fail to accumulate aPKC and Crb properly at their apical membrane. We interpret these results as further evidence of a functional link between Mbt and Cno during polarized photoreceptor morphogenesis.

Our results also show that in *mbt* mutant photoreceptors, Rap1 localization is no longer restricted to the *ZA* but instead spreads apically. Conversely, *Rap1* and *cno* promote the accumulation of Mbt at the *ZA*. Therefore Rap1, Cno and Mbt localization and accumulation at the *ZA* are interlinked. To probe Rap1 function during photoreceptor *ZA* morphogenesis, we assessed the effect of decreasing Rap1 expression on ECad stability. Consistent with the notion that the function of *mbt* and *Rap1* are linked, we find that they are both required to stabilize ECad::GFP at the photoreceptor *ZA.* However, ECad mobile fraction is much higher in *Rap1IR* cells than in *mbt* null cells, evaluated over approximately 1000 sec at approximately 70% for *Rap1IR* and 45% over 250 sec for *mbt^P1^* (Walther et al., 2016). Together with our finding that Mbt accumulation at the *ZA* is decreased in *Rap1IR* cells, our FRAP data are therefore compatible with Mbt mediating part of Rap1’s function in promoting ECad stability. The much larger mobile fraction we estimate in *Rap1IR* when compared to *mbt* null mutant photoreceptors indicates that Rap1 must also regulate ECad stability independently of Mbt. The longer time scale for ECad to recover in *Rap1IR* cells when compared to *mbt* mutant cells is compatible with Rap1 functioning in part through promoting ECad delivery. Previous work in MCF7 cells has shown that Rap1 function in promoting ECad delivery depends on Cdc42 (Hogan et al., 2004). Our work raises the possibility that Pak4 is one of the downstream effector of Cdc42 during this process.

*Rap1* and *cno* have been shown to regulate apical-basal polarity in the cellularizing embryo, a system that allows for examination of the net contribution of the Par complex and *AJ* material toward epithelial cell polarization (Choi et al., 2013). In this model system, *Rap1* and *cno* regulate the apical localization of both Baz and Arm, with Baz reciprocally influencing Cno localization. Later in embryonic development, Baz is required at the *ZA* to capture preassembled *AJ* material, which includes ECad, thus promoting *ZA* morphogenesis (McGill et al., 2009). Our work indicates that similar regulations might be at play during epithelial polarity remodeling. However, unlike in the early embryo, *AJ* material (Arm) is absolutely required for Baz (and Par6-aPKC) accumulation at the cell cortex in the developing pupal photoreceptor (Walther et al., 2016). In this cell, we therefore favor a model whereby Mbt, Cno and Rap1 influence *ZA* morphogenesis primarily through regulating ECad/Arm, through Arm phosphorylation and coupling of ECad/Arm to the actomyosin cytoskeleton and Baz via Cno. Our work shows that in turn these regulations influence apical membrane morphogenesis including aPKC, Part6 and Crb accumulation.

## Acknowledgements

The authors wish to thank all members of the Pichaud lab for helpful discussion, in particular Francisca Nunes Almeida for help with the FRAP assay and for critical reading of the manuscript. We are grateful to Linda Van Aelst and Andreas Wodarz for generously sharing reagents. The N2 A71 monoclonal antibody, developed by Eric Weischaus, was obtained from the Developmental Studies Hybridoma Bank, created by the NICHD of the NIH and maintained at The University of Iowa, Department of Biology, Iowa City, IA 52242. Stocks obtained from the Bloomington Drosophila Stock Center (NIH P40OD018537) were used in this study. This work, including support to RFW, MB and NP was funded by an MRC grant to FP (award code MC_UU_12018/3)

## Material and Methods

**Fly strains:**

The following fly strains were used:

*rap1-Rap1::GFP* and *NP-Gal42631, UAS-Rap1IR* (O’Keefe et al., 2009)

*Rap1IR (BL #29434); bazIR* (*BL #*39072); *cnoIR (BL #33367)*

*dizzy^Δ12^, FRT40A (Huelsmann et al., 2006)*

*dizzy-Dizzy::GFP (Boettner and Van Aelst, 2007)*

*ubi-Cad::GFP* (Oda and Tsukita, 2001)

*mbt^P1^* (Schneeberger and Raabe, 2003)

*mbt^P1^, FRT19A and mbt^P1^, baz^xi106^, FRT9.2 (Walther et al., 2016)*

*w,baz^xi106^, FRT9.2* (Nusslein-Volhard et al., 1987).

*FRT82B, cnoR2 (Sawyer et al., 2009)*

## Analysis of gene function

Clonal analysis of mutant alleles in the retina was performed using the standard FLP-FRT technique (Xu and Rubin, 1993) with appropriate *FRT, ubi-GFP* chromosomes used to generate negatively marked mutant tissue in combination with eyFLP (Newsome et al., 2000). Retina expressing RNAi in clones were generated using the coinFLP system (Bosch et al., 2015). Clones of retinal tissue expressing RNAi against *Rap1* were generated both with and without UAS-dicer, while clones of retinal tissue expressing RNAi against *cno* were generated without UAS-dicer only. In order to mitigate the strong *Rap1* loss of function phenotype, *Rap1IR* animal were raised at 20 degrees and shifted to appropriate temperature (25 or 29 degrees) at puparium formation.

## Antibodies and immunological methods

Whole mount retinas at 40% after puparium formation (APF) were prepared as previously described (Walther and Pichaud, 2006). The following antibodies were used: rabbit anti-PKCζ, 1/600 (SAB4502380, Sigma), mouse anti-Arm, 1/200 (N27-A1, Developmental Studies Hybridoma Bank), rat anti-Baz, 1/1000 (Gift from A.Wodarz, University of Cologne), rabbit anti-Cno, 1/200 (Gift from L. Van Aelst, (Boettner et al., 2003)), rabbit anti-Baz, 1/2000, rat anti-Crb, 1/200, Guinea Pig anti-Mbt 1/200, Guinea Pig anti-PATJ 1/400 (Walther et al., 2016), with the appropriate combination of mouse, guinea pig, rabbit and rat secondary antibodies conjugated to Dy405, Alexa488, Cy3 or Cy5 as appropriate at 1/200 each (Jackson ImmunoResearch) or TRITC-conjugated Phalloidin (P1951, Sigma) at 2μg/mL. Retinas were mounted in VectaShield™ with or without DAPI as appropriate and imaging was performed using a Leica SP5 confocal. Images were edited using ImageJ and Adobe Photoshop 7.0.

## Data analysis

For length and pixel intensity measurements, a threshold was applied to define the *ZA* domain, and a line was drawn along the apical-basal axis of the cell, running in the middle of the *ZA* to measure the length of the Arm, Baz, Mbt domains. Mean pixel intensity was measured using the wand (tracing) tool in Fiji (Schindelin et al., 2012). In all cases, at least four independent mosaic retinas were used for each genotype. To compare the Mbt/Arm or Baz/Arm ratios in individual *ZA*, mean pixel intensity individual measurements for Mbt or Baz and Arm were normalized using the average mean intensity calculated for each epitope for a given set of experiments (i.e Mbt/Arm and Arm/Baz). These normalized data were used to calculate the Mbt/Arm and Baz/Arm ratio for a collection of individual *ZA*. To measure the ratio of apical Rap1::GFP/*ZA* Rap1::GFP, GFP intensity was measured using an ROI covering the *ZA* which was then applied to the apical domain. Statistical analysis was done using Prism 7.0. Data sets were tested for normality (D’Agostino and Pearson normality test) and p-values were calculated using the student’s t-test or the Mann-Whitney test as appropriate.

## Fluorescence recovery after photobleaching (FRAP)

FRAP analysis was performed as previously described (Walther et al., 2016). At 40% APF the pupal cuticle was removed to expose the retina and the animal was mounted in Voltalef oil. Live imaging was performed on a Leica SP5 confocal using a 63x 1.4 NA oil immersion objective at the following settings: pixel resolution 512 x 512, speed 400 Hz, 10% 488 nm laser power at 20% argon laser intensity and 5x zoom. FRAP analysis of *ubi-*ECad::GFP was performed by marking the basal tip of the *AJ* with a 5 pixel-diameter circle ROI followed by photo-bleaching with a single pulse using 90 % 488 nm laser power at 20 % argon laser intensity. *AJ* recovery was recorded every 1.293 seconds with the previously mentioned settings for approximately 1000 sec. FRAP data were drift corrected in Fiji (Schindelin et al., 2012) using the StackReg plugin. Three different z axis profiles were analysed: (1) from the photo-bleached area; (2) from an equivalent area of a neighbouring non-photo-bleached AJ; and (3) from an equivalent area of background. The data were normalized using easyFRAP. ECad::GFP data were fitted to a two-phase association curve in GraphPad Prism. The p values were calculated with an unpaired two-tailed Student’s t test with Welch’s correction.

## Scanning Electron Microscopy

Flies were fixed in 2% paraformaldehyde, 2% glutaraldehyde and 0.1 M cacodylate for 2 hours and then dehydrated in ethanol, as previously described (Richardson and Pichaud, 2010). The samples were then critical-point dried and mounted on aluminum stubs before gold coating. Imaging was carried out on a JEOL Variable Pressure scanning electron microscope (SEM).

---

**Supplementary Figure 1: Rap1 is required to preserve retinal tissue integrity.** (A-A”) *Rap1IR* cells labeled by GFP (A) and stained for Arm (A’). Yellow stars label cone cells in the *Rap 1IR* tissue. White stars label cone cells in one wild type ommatidium. Note the *Rap1IR* ommatidia lack cone cells. A yellow dashed box highlights *Rap1IR* ommatidia lacking interommatidial cells. (B-D) *Rap1IR* cell labeled by GFP (blue) and stained for Arm (B’, C’, D’) aPKC (B”, C”, D”) and Mbt (B”’, C”’, D”’). Note that many *Rap1IR* photoreceptors delaminate below the floor of the retina, white arrows (D-D””). Scale bars = 2μm.

**Supplementary Figure 2: Genetic modifiers of the *Rap1IR* rough eye phenotype.** (A) SEM of a wild type eye, (B) *Rap1IR*, (C) Heterozygous *dizzydΔ^12^* eye, (D) *Rap1IR* combined with *dizzydΔ^12^* / +, (E) *bazIR,* (F) *bazIR* combined with *RapIR.*

**Supplementary Figure 3: Mbt promotes Arm and Baz accumulation at the *ZA***. (A-B) *mbt^P1^* mutant cells labeled by the lack of nuclear GFP (blue) and stained for Baz (A’ and B”) and Crb (A”). Scale bars = 2μ.

## REFERENCES

Berger, S., Bulgakova, N. A., Grawe, F., Johnson, K. and Knust, E. (2007). Unraveling the genetic complexity of Drosophila stardust during photoreceptor morphogenesis and prevention of light-induced degeneration. Genetics 176, 2189–2200.

Boettner, B., Harjes, P., Ishimaru, S., Heke, M., Fan, H. Q., Qin, Y., Van Aelst, L. and Gaul, U. (2003). The AF-6 homolog canoe acts as a Rap1 effector during dorsal closure of the Drosophila embryo. Genetics 165, 159–169.

Boettner, B. and Van Aelst, L. (2007). The Rap GTPase activator Drosophila PDZ-GEF regulates cell shape in epithelial migration and morphogenesis. Mol Cell Biol 27, 7966–7980.

Bosch, J. A., Tran, N. H. and Hariharan, I. K. (2015). CoinFLP: a system for efficient mosaic screening and for visualizing clonal boundaries in Drosophila. Development 142, 597–606.

Bryant, D. M. and Stow, J. L. (2004). The ins and outs of E-cadherin trafficking. Trends Cell Biol 14, 427–434.

Choi, W., Harris, N. J., Sumigray, K. D. and Peifer, M. (2013). Rap1 and Canoe/afadin are essential for establishment of apical-basal polarity in the Drosophila embryo. Mol Biol Cell 24, 945–963.

Hogan, C., Serpente, N., Cogram, P., Hosking, C. R., Bialucha, C. U., Feller, S. M., Braga, V. M., Birchmeier, W. and Fujita, Y. (2004). Rap1 regulates the formation of E-cadherin-based cell-cell contacts. Mol Cell Biol 24, 6690–6700.

Hong, Y., Ackerman, L., Jan, L. Y. and Jan, Y. N. (2003). Distinct roles of Bazooka and Stardust in the specification of Drosophila photoreceptor membrane architecture. Proc Natl Acad Sci USA 100, 12712–12717.

Huelsmann, S., Hepper, C., Marchese, D., Knoll, C. and Reuter, R. (2006). The PDZ-GEF dizzy regulates cell shape of migrating macrophages via Rap1 and integrins in the Drosophila embryo. Development 133, 2915–2924.

Izaddoost, S., Nam, S. C., Bhat, M. A., Bellen, H. J. and Choi, K. W. (2002). Drosophila Crumbs is a positional cue in photoreceptor adherens junctions and rhabdomeres. Nature 416, 178–183.

Jin, D., Durgan, J. and Hall, A. (2015). Functional cross-talk between Cdc42 and two downstream targets, Par6B and PAK4. Biochem J 467, 293–302.

Knox, A. L. and Brown, N. H. (2002). Rap1 GTPase regulation of adherens junction positioning and cell adhesion. Science 295, 1285–1288.

Kooistra, M. R., Dube, N. and Bos, J. L. (2007). Rap1: a key regulator in cell-cell junction formation. J Cell Sci 120, 17–22.

Krahn, M. P., Buckers, J., Kastrup, L. and Wodarz, A. (2010). Formation of a Bazooka-Stardust complex is essential for plasma membrane polarity in epithelia. J Cell Biol 190, 751–760.

Law, S. H. and Sargent, T. D. (2014). The serine-threonine protein kinase PAK4 is dispensable in zebrafish: identification of a morpholino-generated pseudophenotype. PLoS One 9, e100268.

Mandai, K., Rikitake, Y., Shimono, Y. and Takai, Y. (2013). Afadin/AF-6 and canoe: roles in cell adhesion and beyond. Prog Mol Biol Transl Sci 116, 433–454.

McGill, M. A., McKinley, R. F. and Harris, T. J. (2009). Independent cadherin-catenin and Bazooka clusters interact to assemble adherens junctions. J Cell Biol 185, 787–796.

Menzel, N., Melzer, J., Waschke, J., Lenz, C., Wecklein, H., Lochnit, G., Drenckhahn, D. and Raabe, T. (2008). The Drosophila p21-activated kinase Mbt modulates DE-cadherin-mediated cell adhesion by phosphorylation of Armadillo. Biochem J 416, 231–241.

Menzel, N., Schneeberger, D. and Raabe, T. (2007). The Drosophila p21 activated kinase Mbt regulates the actin cytoskeleton and adherens junctions to control photoreceptor cell morphogenesis. Mech Dev 124, 78–90.

Morais-de-Sa, E., Mirouse, V. and St Johnston, D. (2010). aPKC phosphorylation of Bazooka defines the apical/lateral border in Drosophila epithelial cells. Cell 141,509–523.

Nam, S. C. and Choi, K. W. (2003). Interaction of Par-6 and Crumbs complexes is essential for photoreceptor morphogenesis in Drosophila. Development 130,4363–4372.

Newsome, T. P., Asling, B. and Dickson, B. J. (2000). Analysis of Drosophila photoreceptor axon guidance in eye-specific mosaics. Development 127, 851–860.

Nusslein-Volhard, C., Frohnhofer, H. G. and Lehmann, R. (1987). Determination of anteroposterior polarity in Drosophila. Science 238, 1675–1681.

O’Keefe, D. D., Gonzalez-Nino, E., Burnett, M., Dylla, L., Lambeth, S. M., Licon, E., Amesoli, C., Edgar, B. A. and Curtiss, J. (2009). Rap1 maintains adhesion between cells to affect Egfr signaling and planar cell polarity in Drosophila. Dev Biol 333,143–160.

Oda, H. and Tsukita, S. (2001). Real-time imaging of cell-cell adherens junctions reveals that Drosophila mesoderm invagination begins with two phases of apical constriction of cells. J Cell Sci 114, 493–501.

Pellikka, M., Tanentzapf, G., Pinto, M., Smith, C., McGlade, C. J., Ready, D. F. and Tepass, U. (2002). Crumbs, the Drosophila homologue of human CRB1/RP12, is essential for photoreceptor morphogenesis. Nature 416, 143–149.

Ready, D. F. (2002). Drosophila compound eye morphogenesis: Blind mechanical engineers? In Results and Problems in Cell Differentiation (ed. K. Moses), pp. 191–204. Berlin Heildelderg New York: Springer-Verlag.

Richard, M., Grawe, F. and Knust, E. (2006). DPATJ plays a role in retinal morphogenesis and protects against light-dependent degeneration of photoreceptor cells in the Drosophila eye. Dev Dyn 235, 895–907.

Richardson, E. C. and Pichaud, F. (2010). Crumbs is required to achieve proper organ size control during Drosophila head development. Development 137, 641–650.

Sawyer, J. K., Harris, N. J., Slep, K. C., Gaul, U. and Peifer, M. (2009). The Drosophila afadin homologue Canoe regulates linkage of the actin cytoskeleton to adherens junctions during apical constriction. J Cell Biol 186,57–73.

Schindelin, J., Arganda-Carreras, I., Frise, E., Kaynig, V., Longair, M., Pietzsch, T., Preibisch, S., Rueden, C., Saalfeld, S., Schmid, B., et al. (2012). Fiji: an open-source platform for biological-image analysis https://sites.google.com/site/qingzongtseng/imagejplugins. Nature methods 9, 676–682.

Schneeberger, D. and Raabe, T. (2003). Mbt, a Drosophila PAK protein, combines with Cdc42 to regulate photoreceptor cell morphogenesis. Development 130,427–437.

Spahn, P., Ott, A. and Reuter, R. (2012). The PDZ-GEF protein Dizzy regulates the establishment of adherens junctions required for ventral furrow formation in Drosophila. J Cell Sci 125, 3801–3812.

Tepass, U. (2012). The apical polarity protein network in Drosophila epithelial cells: regulation of polarity, junctions, morphogenesis, cell growth, and survival. Annu Rev Cell Dev Biol 28, 655–685.

Wallace, S. W., Durgan, J., Jin, D. and Hall, A. (2010). Cdc42 regulates apical junction formation in human bronchial epithelial cells through PAK4 and Par6B. Mol Biol Cell 21, 2996–3006.

Walther, R. F., Nunes de Almeida, F., Vlassaks, E., Burden, J. J. and Pichaud, F. (2016). Pak4 Is Required during Epithelial Polarity Remodeling through Regulating AJ Stability and Bazooka Retention at the ZA. Cell reports 15, 45–53.

Walther, R. F. and Pichaud, F. (2006). Immunofluorescent staining and imaging of the pupal and adult Drosophila visual system. Nat Protoc 1, 2635–2642.

Walther, R. F. and Pichaud, F. (2010). Crumbs/DaPKC-dependent apical exclusion of Bazooka promotes photoreceptor polarity remodeling. Curr Biol 20, 1065–1074.

Wang, Y. C., Khan, Z. and Wieschaus, E. F. (2013). Distinct Rap1 activity states control the extent of epithelial invagination via alpha-catenin. Dev Cell 25, 299–309.

Wei, S. Y., Escudero, L. M., Yu, F., Chang, L. H., Chen, L. Y., Ho, Y. H., Lin, C. M., Chou, C. S., Chia, W., Modolell, J., et al. (2005). Echinoid is a component of adherens junctions that cooperates with DE-Cadherin to mediate cell adhesion. Dev Cell 8, 493–504.

Xu, T. and Rubin, G. M. (1993). Analysis of genetic mosaics in developing and adult Drosophila tissues. Development 117, 1223–1237.

